# Subthalamic nucleus activity and its pallidal and nigral outputs shape motor control

**DOI:** 10.64898/2026.11.25.769089

**Authors:** Tom Macpherson, Tianchang Zhao, Monica Natsumi Daudelin, Tadaaki Nishioka, Suthinee Attachaipanich, Takaaki Ozawa, Takatoshi Hikida

## Abstract

The subthalamic nucleus (STN) contributes to motor control, but how its different output pathways regulate spontaneous movement and motor task performance remains unclear. Although classical basal ganglia models emphasize movement suppression by the STN, its influence on different aspects of motor behavior requires further experimental investigation. Here, we examined how manipulation of STN neurons and their projections to the substantia nigra pars reticulata (SNr) and external globus pallidus (GPe) affects spontaneous activity and rotarod performance. We combined optogenetic excitation and inhibition with anatomical tracing in Pitx2-Cre mice. Somatic excitation reduced spontaneous mobility and induced jumping, whereas inhibition increased mobility. Both excitation and inhibition disrupted rotarod performance, suggesting that successful task performance depends on maintaining STN activity within an appropriate range. Anatomical tracing revealed overlapping projections to the SNr and GPe, and excitation of STN axons within either region reduced mobility and disrupted rotarod performance. SNr-targeted inhibition increased spontaneous movement without a detectable change in rotarod performance. These findings support a movement-suppressing role for the STN while highlighting the importance of appropriately regulated STN activity for motor task performance. They identify both nigral and pallidal projections as routes through which STN activation can alter motor behavior.

## 1. Introduction

Successful movement requires the coordination of action initiation, ongoing execution, and postural support. Basal ganglia outputs influence these processes through thalamic and brainstem circuits, including pathways that regulate locomotion and muscle tone (Freeze et al., 2013; Roseberry et al., 2016; Takakusaki et al., 2003; Macpherson et al., 2021). Within this network, the subthalamic nucleus (STN) provides excitatory input to pallidal and nigral structures (Kita and Kitai, 1987; Nambu et al., 2000). In classical models, recruitment of the STN through the indirect pathway strengthens inhibitory basal ganglia output, restricting movement, whereas the direct pathway facilitates movement by reducing this output (Nambu et al., 2002). Although direct and indirect pathways can exert opposing effects on movement (Freeze et al., 2013; Kravitz et al., 2010), their coordinated recruitment also contributes to action selection and initiation (Cui et al., 2013; Macpherson et al., 2014; Nishioka et al., 2023). Understanding the STN’s contribution to motor control therefore requires examining how its activity and downstream projections influence both spontaneous movement and ongoing motor task performance.

The STN also receives motor cortical input through the hyperdirect pathway, which bypasses the striatum and rapidly conveys excitation to basal ganglia output nuclei (Nambu et al., 2000; Nambu et al., 2002). Physiological experiments distinguish early hyperdirect excitation from later excitation transmitted through the striatum and GPe (Polyakova et al., 2020). Eliminating cortico-STN projections increases spontaneous locomotion in mice (Koketsu et al., 2021), while disrupting striatopallidal transmission reduces late nigral excitation and produces hyperactivity (Sano et al., 2013). These findings support a role for temporally organized signaling through the STN in regulating when movements occur. Consistent with this view, STN neurons respond during action switching and cancellation as well as movement preparation and execution (Isoda and Hikosaka, 2008; Polyakova et al., 2024; Schmidt et al., 2013).

Evidence concerning the STN itself spans behavioral interruption, movement generation, and the maintenance of motor performance. Brief STN excitation can interrupt ongoing licking (Fife et al., 2017). In freely moving mice, somatic excitation reduces locomotor measures while inducing grooming and jumping, and disrupts performance on rotarod and balance-beam tasks (Guillaumin et al., 2021). Conversely, quantitative kinematic analysis has shown that STN stimulation can generate rapid head and body movements through brainstem projections (Friedman and Yin, 2023). Excitation of parafascicular thalamic projections to the STN—the “super-direct” pathway—can initiate movement and alleviate akinesia in dopamine-depleted mice (Watson et al., 2021). Movement-related STN activity also supports ongoing locomotion (Callahan et al., 2024), and reducing STN activity in monkeys increases pallidal spike-train variability and impairs movement execution (Hasegawa et al., 2022). Together, these findings suggest that the effect of STN perturbation depends on the recruited circuit, temporal pattern, and behavioral measure.

The STN is heterogeneous at both the molecular and circuit levels. Spatially patterned molecular expression and overlapping genetically defined populations have been identified within the mouse STN (Parolari et al., 2021; Wallén-Mackenzie et al., 2020), and motor cortical projections form somatotopically organized territories in primates (Nambu et al., 1996). STN axons innervate multiple downstream targets and can branch to reach more than one structure (Kita and Kitai, 1987; Sato et al., 2000). Two prominent targets are the substantia nigra pars reticulata (SNr) and external globus pallidus (GPe) (Kita and Kitai, 1987). The SNr is a major inhibitory output nucleus (Freeze et al., 2013), whereas the GPe contains interconnected cell populations with different projection patterns and responses to striatal and STN inputs (Abdi et al., 2015; Aristieta et al., 2021; Mallet et al., 2012). These arrangements could allow STN output to produce different or interacting effects across its targets. Previous studies have linked molecularly defined STN populations to distinct behavioral functions and shown that manipulating STN projections can alter locomotion and movement patterns (Parolari et al., 2021; Ji et al., 2023; Zhou et al., 2026). However, these findings leave unresolved how the STN’s influence on spontaneous movement relates to its contribution to maintaining ongoing motor task performance. In particular, whether its nigral and pallidal outputs contribute similarly to these two aspects of motor control, and how their contributions relate to the effects of excitation and inhibition at the level of STN cell bodies, remain unclear.

Here, we examined how STN neurons and their projections to the SNr and GPe regulate spontaneous movement and motor task performance. We combined optogenetic excitation and inhibition of Pitx2-Cre-targeted STN neurons and their axons with open-field and rotarod testing, alongside anatomical tracing to characterize overlap between the two projection populations. Comparing somatic and axonal manipulations allowed us to assess which behavioral effects of altering STN activity were reproduced by manipulating its outputs within each target.

## 2. Materials and Methods

### 2.1. Animals

Heterozygous Pitx2-Cre mice (129S-Pitx2^tm4(cre)Jfm/Mmucd; MMRRC; RRID:MMRRC_000126-UCD) maintained on a C57BL/6N background and aged 10-20 weeks were used for all experiments. Males and females were counterbalanced across conditions and data were grouped as no significant main effect or interactions of sex were observed. Animals were housed in groups of two to four under a 12-hour light/dark cycle with lights on at 08:00. Room temperature was approximately 24 ± 2 °C and humidity 50 ± 5%, with food and water available ad libitum. All procedures conformed to the guidelines of the National Institutes of Health experimental procedures and were approved by the Ethical Committee of the Institute for Protein Research at Osaka University.

### 2.2. Stereotaxic virus infusion and optic fiber implantation

Mice were anesthetized with ketamine (90 mg/kg) and xylazine (20 mg/kg), administered intraperitoneally, and positioned in a stereotaxic apparatus (Stoelting co.). A midline scalp incision was made, followed by a craniotomy using a dental drill. Viral injections were made through graduated pipettes with tip diameters of 10–15 μm.

#### 2.2.1. Optogenetic experiments

Mice were infused bilaterally into the STN (bregma coordinates: AP, −1.90 mm; ML, ± 1.50 mm; DV, −4.50 mm and −4.25 mm from brain surface, 250 nl/site at a rate of 100 nl/min) with a Cre-dependent adeno-associated virus (AAV) expressing channelrhodopsin-2 (AAV5-Ef1a-DIO-hChR2(H134R)-EYFP-WPRE-HGHpA; catalog #20298, Addgene) at a concentration of ≥ 1×10¹³ vg/mL, archaerhodopsin (AAV5-CAG-FLEX-ArchT-tdTomato; catalog #28305, Addgene) at a concentration of ≥ 7×10¹² vg/mL, JAWS (AAV5-CAG-FLEX-rc [Jaws-KGC-GFP-ER2]; #84445, Addgene) at a concentration of ≥ 7×10¹² vg/mL, or an optically inactive control virus (AAV5-Ef1a-DIO-EYFP; catalog #27056, Addgene) at a concentration of ≥ 1×10¹³ vg/mL, using a Nanoject III injection system (Drummond Instruments). Following infusion, the needle was kept at the injection site for 5 min to allow for diffusion of the virus and then slowly withdrawn to reduce backflow. Two weeks after viral infusion, mice were implanted with optic fibers (core, 200 μm; 0.22 numerical aperture; Thorlabs, NJ, USA) threaded through ceramic zirconia ferrules into the STN (bregma coordinates: AP, −1.90 mm; ML, ± 1.50 mm; DV, −4.25 mm from brain surface), SNr (bregma coordinates: AP, -3.30 mm; ML, ± 1.00 mm; DV, −4.50mm from brain surface), or GPe (bregma coordinates: AP, −0.50 mm; ML, ± 1.75 mm; DV, −3.00 mm from brain surface). Cranial screws were implanted 1 mm into the skull surrounding the optic fibers, and optic fibers were secured to the skull using dental cement. Following optic fiber implantation, mice were left in their home cage for a further 2-3 weeks to ensure recovery from surgery and further viral expression prior to experiments commencing.

#### 2.2.2. Anterograde tracing of neuronal processes and synaptophysin-associated presynaptic structures

Mice were infused unilaterally into the STN (bregma coordinates: AP, −1.90 mm; ML, ± 1.50 mm; DV, −4.50 mm and −4.25 mm from brain surface, 250 nl/site at a rate of 100 nl/min) with AAVDJ-phSyn(s)-FLEX-tdTomato-T2A-sypEGFP-WPRE (catalog #51509, Addgene) at a concentration of ≥ 4×10¹² vg/mL. The needle remained in place for 5 minutes after injection. Tissue was collected after three weeks of expression.

#### 2.2.3. Dual retrograde tracing

Mice were infused unilaterally on the same side into the SNr (bregma coordinates: AP −3.40 mm, ML ±1.40 mm, and DV −4.25 mm from brain surface, 250 nl/site at a rate of 100 nl/min) and GPe (AP−0.50 mm, ML ±1.75 mm, and DV −3.00/−3.75 mm, with 250 nl per site at a rate of 100 nl/min) with AAVretro-hSyn-DIO-EGFP (#50457, Addgene) and AAVretro-hSyn-DIO-mCherry (#50459, Addgene), with one fluorophore assigned to each target and assignments counterbalanced across mice. The needle remained in place for 5 minutes after injection. Tissue was collected after five weeks of expression.

### 2.3. Histology and quantification

Under ketamine (90 mg/kg) and xylazine (20 mg/kg) anesthesia, mice were transcardially perfused with 0.1 M PBS for 2 min, followed by 4% paraformaldehyde (PFA; Nacalai Tesque) in 1× PBS for 5 min at a flow rate of 10 ml/min. Brains were removed, postfixed overnight in 4% PFA, and sequentially cryoprotected in 7.5%, 15%, and 30% sucrose in 1× PBS at 4°C, allowing the tissue to sink at each stage. Brains were then embedded in Optimal Cutting Temperature (O.C.T.) compound (Sakura Finetek), frozen, and mounted on a cryostat specimen holder. Coronal sections (40 μm thick) were cut using a cryostat (CM1860; Leica) at −17 to −20°C.

For Fos immunostaining, free-floating sections were permeabilized in 0.25% Triton X-100 in PBS for 5 min at room temperature and blocked in 5% normal goat serum in PBS (pH 7.4) for 1 h at room temperature with gentle agitation. Sections were incubated with rabbit monoclonal anti-c-Fos antibody (1:100; clone EPR20769; Abcam, ab214672) overnight at 4°C, followed by Alexa Fluor 568-conjugated goat anti-rabbit IgG (H+L) cross-adsorbed secondary antibody (1:1000; Invitrogen, A-11011) for 2 h. Sections were washed three times in PBS for 1 min each after permeabilization, blocking, and each antibody incubation.

Sections were mounted on slides and coverslipped using Fluoroshield mounting medium containing DAPI (Abcam). Fluorescence images were acquired using a BZ-X800E All-in-One Fluorescence Microscope (KEYENCE). Retrogradely labeled cells were quantified using HALO software (Indica Labs). DAPI-labeled, EGFP-labeled, mCherry-labeled, and EGFP- and mCherry double-labeled cells were classified within the STN, and counts were expressed relative to the number of DAPI-positive cells.

### 2.4. Optogenetic Procedures

#### 2.4.1. Somatic optogenetic experiments

Mice received 473 nm laser stimulation at 5Hz, 20 Hz, and 40Hz for ChR2-expressing or EYFP-expressing mice, or 532 nm at a constant rate for ArchT-expressing mice or EYFP-expressing mice. The same mice underwent 5-, 20-, and 40-Hz stimulation on consecutive days in a Latin-square design. This frequency-order design was used for both the open-field and rotarod experiments.

#### 2.4.2. Axon terminal optogenetic experiments

Mice received 473 nm laser stimulation at 20 Hz for ChR2-expressing or EYFP-expressing mice, or 638 nm at a constant rate for JAWS-expressing mice or EYFP-expressing mice. Laser stimulation was delivered by DPSS (diode-pumped solid-state) lasers (Shanghai Laser & Optics Century Co., Ltd.) controlled by a microcontroller (Arduino) at an intensity of 5 mW at the fiber tip for somatic stimulation and 10 mW at the fiber tip for axon terminal stimulation.

### 2.5. Behavioral Testing

#### 2.5.1. Open field test (OFT)

Mice were placed in a gray cylindrical open-field arena (42 cm in diameter and 42 cm in height). Patch cords were connected to bilateral fiber-optic cannulae and suspended above the mice to allow unrestricted movement throughout the arena. Following a 3-min habituation period, mice underwent a 12-min test comprising four alternating 3-min laser OFF and ON periods (OFF–ON–OFF–ON), as previously described (Attachaipanich et al., 2023; Tye et al., 2013). This design enabled assessment of locomotor responses during illumination and any persisting effects during the subsequent laser OFF period. Photostimulation was delivered using the parameters described in 2.4. Percentage of time spent mobile (Time Mobile %), distance travelled (cm), and velocity (cm/s) were quantified automatically using video-tracking software (EthoVision XT 16; Noldus). Mobility in “Time Mobile %” was classified using velocity thresholds of 2.0 cm/s for movement onset and 1.75 cm/s for movement cessation. Jumping during the OFT was quantified manually by observation of videos captured from above the arena. Jumping was defined as a discrete movement in which the mouse propelled its body off the arena floor, with all four paws simultaneously losing contact with the floor.

#### 2.5.2. Rotarod test

Mice were tested for their ability to stay on a rotating bar (rotarod, Harvard Apparatus), which was kept at a constant rotation of 2 rpm. For each mouse, two trials were performed. First a trial in which the laser was switched off (laser OFF), then, immediately following, a trial in which the laser was constantly on (laser ON). Latency to fall (in seconds) was scored for each trial, with trials ending after a fall or a maximum of 120 seconds.

### 2.6. Statistical analysis

In the OFT, for each mouse and session, percentage of time spent mobile and mean velocity were calculated for each 3-min period, whereas distance travelled was calculated as the total distance covered during that period. These values were then averaged across the two laser OFF periods and separately across the two laser ON periods. Percentage of time spent mobile, distance travelled, velocity, jump count, and rotarod latency were analyzed separately using two-way repeated measures ANOVAs, with virus (ChR2, ArchT, or Jaws versus the corresponding eYFP control) as the between-subjects factor and laser condition (OFF versus ON) as the within-subjects factor. Following ANOVA, pairwise comparisons were performed between laser conditions within each virus group and between virus groups within each laser condition using Fisher’s least significant difference (LSD) tests. Each comparison family contained a single comparison; therefore, no multiplicity adjustment was applied. For dual retrograde tracing, percentages of SNr-only, GPe-only, and double-labeled cells were compared using a one-way repeated-measures ANOVA, with labeling category as the within-subjects factor and the Greenhouse–Geisser correction, followed by Tukey’s multiple-comparisons test. Residual Q–Q plots and residual-versus-fitted plots were visually inspected; no apparent outliers or substantial differences in variability between groups were identified. Jump counts were not analyzed when all observations were zero. Statistical analyses were performed using GraphPad Prism v11. Complete ANOVA results and pairwise comparisons are reported in Supplementary Table 1.

## 3. Results

### 3.1. Optogenetic targeting of STN neurons in Pitx2-Cre mice

To examine how STN activity influences spontaneous movement and motor task performance, we used Pitx2-Cre mice to direct opsin expression to STN neurons. Pitx2 encodes paired-like homeodomain transcription factor 2, which is required for normal STN development and is broadly expressed in glutamatergic neurons throughout the adult STN (Martin et al., 2004; Schweizer et al., 2016). Building on previous genetic targeting of Pitx2-expressing STN neurons (Schweizer et al., 2014), we combined this Cre-driver line with local delivery of Cre-dependent viral vectors to preferentially target STN neurons over neighboring structures (Nishioka et al., 2020).

For somatic manipulation, Cre-dependent adeno-associated viral vectors encoding either the excitatory channelrhodopsin ChR2 or the inhibitory proton pump ArchT were injected bilaterally into the STN, with separate control groups receiving a vector encoding eYFP alone. Optical fibers were implanted bilaterally above the STN to enable illumination during behavioral testing (Fig. 1A,B). Following photostimulation, Fos labeling was observed in ChR2-expressing STN neurons, supporting the effectiveness of this approach in stimulating STN neuronal activity (Fig. 1B).

**Figure 1.**
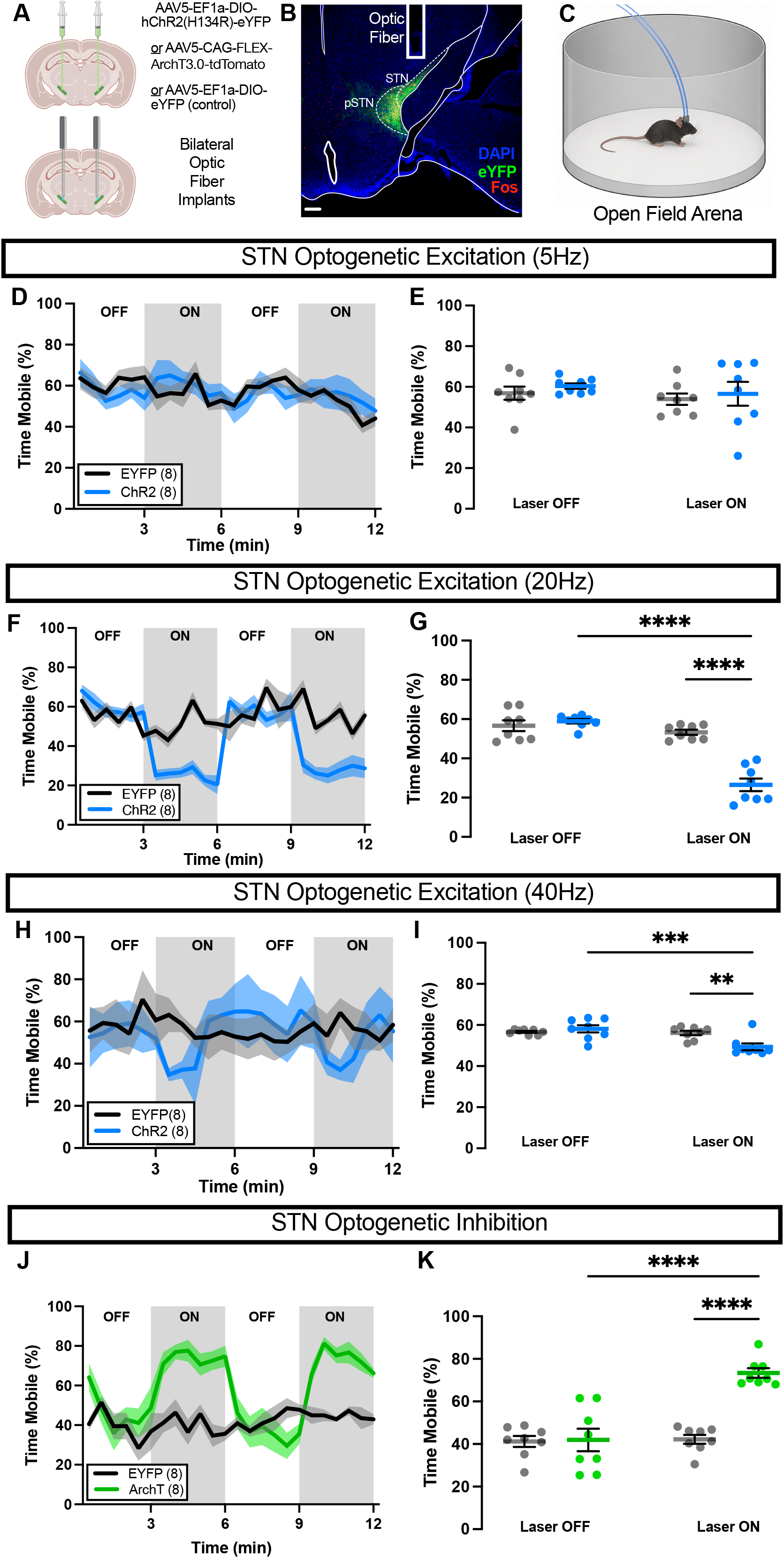
Somatic STN excitation and inhibition have opposing effects on open-field mobility. (A) Bilateral Cre-dependent viral delivery and optical fiber implantation in the STN of Pitx2-Cre mice. (B) Representative eYFP expression, Fos labeling, DAPI counterstaining, and fiber location. (C) Open-field apparatus. (D,E), (F,G), and (H,I) Mobility time courses and OFF/ON summaries during ChR2 excitation at 5, 20, and 40 Hz, respectively. (J,K) Corresponding measures during continuous ArchT inhibition. Each opsin cohort had a separate eYFP control cohort; the same excitation mice underwent all three frequencies. Gray shading denotes illumination periods. Data are shown as individual mice and group means ± SEM; *n* = 8 mice per virus group. Two-way repeated measures ANOVA followed by unadjusted Fisher’s LSD comparisons. Asterisks denote the bracketed pairwise comparisons: \**p* < 0.05, \*\**p* < 0.01, \*\*\**p* < 0.001, \*\*\*\**p* < 0.0001. Scale bars in B: 200 μm. STN: subthalamic nucleus, pSTN: posterior subthalamic nucleus. Diagrams in A and C generated using BioRender.com.

### 3.2. STN excitation and inhibition bidirectionally regulate spontaneous movement

The effects of STN excitation on spontaneous movement were assessed in an open field during alternating laser OFF and ON periods. ChR2-expressing mice and their eYFP controls underwent separate sessions with stimulation at 5, 20, and 40 Hz. Stimulation at 5 Hz did not significantly alter mobile time (Fig. 1D,E), whereas stimulation at 20 Hz (virus × laser interaction, *F*(1, 14) = 94.94, *p* < 0.0001; Fig. 1F,G) and 40 Hz (*F*(1, 14) = 8.569, *p* = 0.0110; Fig. 1H,I) reduced mobility. The largest observed reduction occurred at 20 Hz.

Despite reduced mobile time, distance travelled and measured velocity increased during 20-Hz stimulation (virus × laser interactions: distance, *F*(1, 14) = 22.80, *p* = 0.0003; velocity, *F*(1, 14) = 20.20, *p* = 0.0005; Supplementary Fig. 1D,E). Velocity also increased at 40 Hz (*F*(1, 14) = 5.062, *p* = 0.0411; Supplementary Fig. 1H). The increase in distance at 40 Hz was supported by pairwise comparisons, although the interaction was nonsignificant (*F*(1, 14) = 4.079, *p* = 0.0630; Supplementary Fig. 1G). Behavioral recordings revealed sporadic jumps across the arena and into its walls. Jumping increased at both 20 Hz (*F*(1, 14) = 33.83, *p* < 0.0001) and 40 Hz (*F*(1, 14) = 15.56, *p* = 0.0015; Supplementary Fig. 1F,I), suggesting that these abrupt displacements largely accounted for the elevated distance and velocity.

In contrast, inhibition of STN neurons increased spontaneous mobility. ArchT-expressing mice showed increases in the percentage of time spent mobile (virus × laser interaction, *F*(1, 14) = 42.32, *p* < 0.0001; Fig. 1J,K), distance travelled (*F*(1, 14) = 116.1, *p* < 0.0001; Supplementary Fig. 1J), and velocity (*F*(1, 14) = 123.4, *p* < 0.0001; Supplementary Fig. 1K). No jumping was observed in either group (Supplementary Fig. 1L). Thus, excitation and inhibition of STN neurons had opposing effects on open-field movement, with jumping specifically accompanying excitation.

### 3.3. Both excitation and inhibition of STN neurons disrupt rotarod performance

We next examined how somatic STN manipulation affected performance on a slowly rotating rod, a task requiring coordinated stepping and postural adjustments (Fig. 2A). Stimulation at 5 Hz did not significantly alter latency to fall (Fig. 2B). In contrast, stimulation at 20 Hz (virus × laser interaction, *F*(1, 14) = 31,223, *p* < 0.0001; Fig. 2C) and 40 Hz (*F*(1, 14) = 8.844, *p* = 0.0101; Fig. 2D) shortened the fall latency. The greatest observed reduction occurred at 20 Hz, when mice jumped off the rod immediately after light onset, indicating that the reduced fall latency could not be attributed specifically to impaired balance.

**Figure 2.**
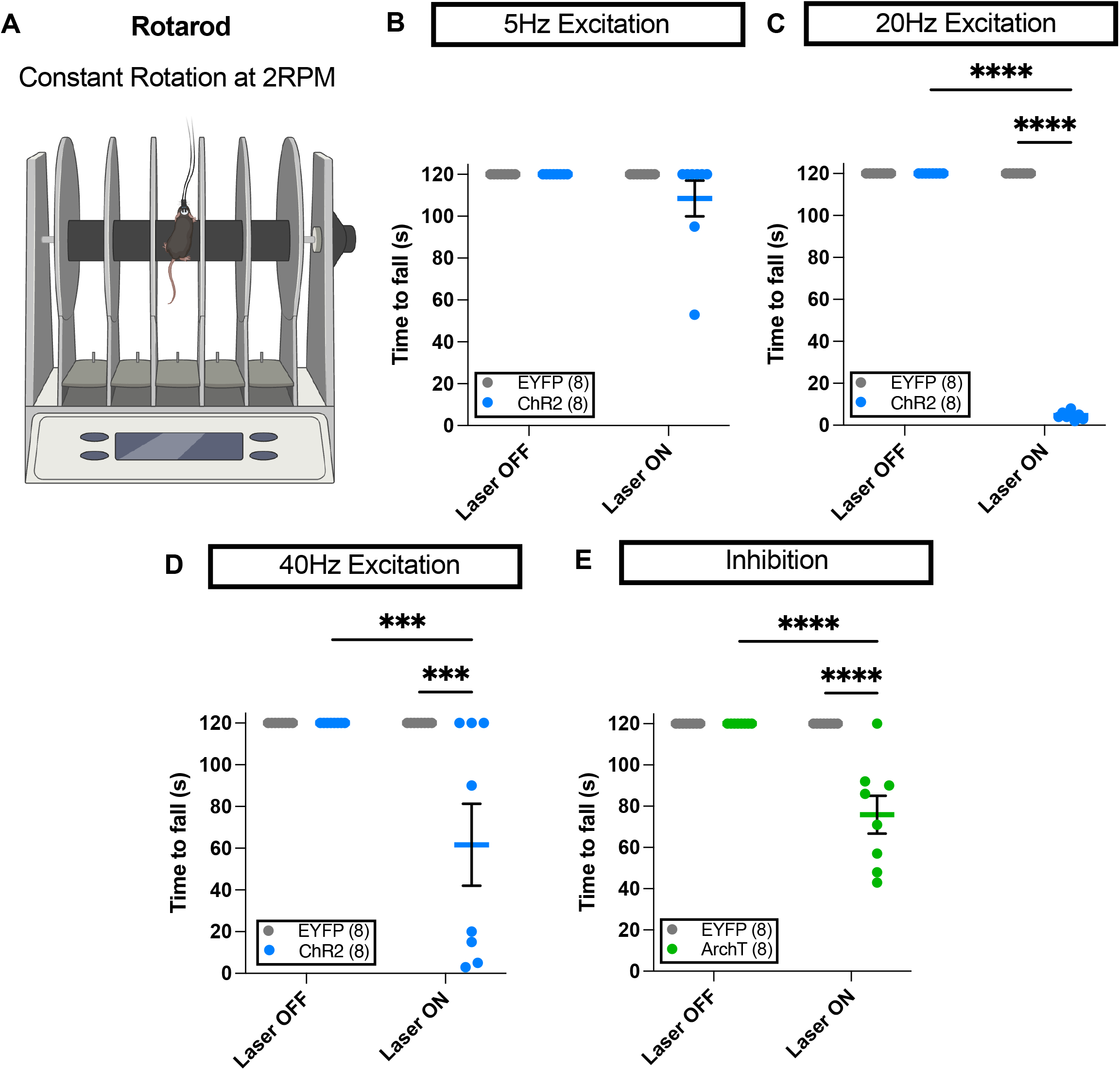
Somatic STN excitation and inhibition shorten rotarod latency. (A) Schematic of rotarod experimental setup (diagram was generated using BioRender.com). (B–D) Latency to fall during OFF and ON trials in ChR2-expressing mice and controls at 5, 20, and 40 Hz. (E) Corresponding ArchT inhibition experiment. Trials ended at a fall or 120 s. Data are shown as individual mice and group means ± SEM; *n* = 8 mice per virus group. Two-way repeated measures ANOVA followed by unadjusted Fisher’s LSD comparisons. Asterisks denote the bracketed pairwise comparisons: \**p* < 0.05, \*\**p* < 0.01, \*\*\**p* < 0.001, \*\*\*\**p* < 0.0001.

Despite increasing spontaneous movement in the open field, STN inhibition also shortened rotarod fall latency (virus × laser interaction, *F*(1, 14) = 22.97, *p* = 0.0003; Fig. 2E). ArchT-expressing mice were observed to run forwards and fall from the rod during illumination. Although these behavioral responses were not systematically quantified, they suggest that excitation and inhibition disrupted task performance in different ways. The impairment following both manipulations suggests that successful task performance depends on maintaining STN activity within an appropriate range.

### 3.4. Overlapping populations of STN neurons project to the SNr and GPe

Somatic manipulation established that changes in STN activity affect both spontaneous movement and rotarod performance, but did not identify the output pathways contributing to these effects. We therefore focused on the substantia nigra pars reticulata (SNr) and external globus pallidus (GPe), two major STN targets that occupy different positions within basal ganglia motor circuits. The SNr is a principal basal ganglia output nucleus, whereas the GPe regulates activity within the basal ganglia and forms reciprocal connections with the STN (Abdi et al., 2015; Aristieta et al., 2021; Freeze et al., 2013; Kita and Kitai, 1987; Mallet et al., 2012). Before examining their behavioral contributions, we used Cre-dependent anterograde and retrograde viral tracing in Pitx2-Cre mice to characterize these projections and determine the extent to which their neurons of origin overlap.

Anterograde tracing following injection of a Cre-dependent viral vector into the STN (Fig. 3A,B) revealed tdTomato-labeled axons and synaptophysin-positive puncta (EGFP) in both the SNr (Fig. 3C) and GPe (Fig. 3D), consistent with the presence of STN-derived presynaptic terminals in these regions. Having confirmed innervation of both targets, we next asked whether these projections originate from separate or overlapping populations of STN neurons. Cre-dependent retrograde viral vectors encoding distinct fluorescent reporters were injected into the SNr and GPe of the same mice (Fig. 3E-G). This approach identified STN neurons labeled from either target alone, as well as neurons expressing both reporters (Fig. 3H, H’). Approximately 25% of DAPI-labeled cells within the STN were positive for both reporters, demonstrating overlapping projections to the SNr and GPe among Pitx2-Cre-targeted STN neurons. Smaller populations were labeled only by the tracer injected into the SNr or GPe, although percentages did not differ significantly across labeling categories (Fig. 3I).

**Figure 3.**
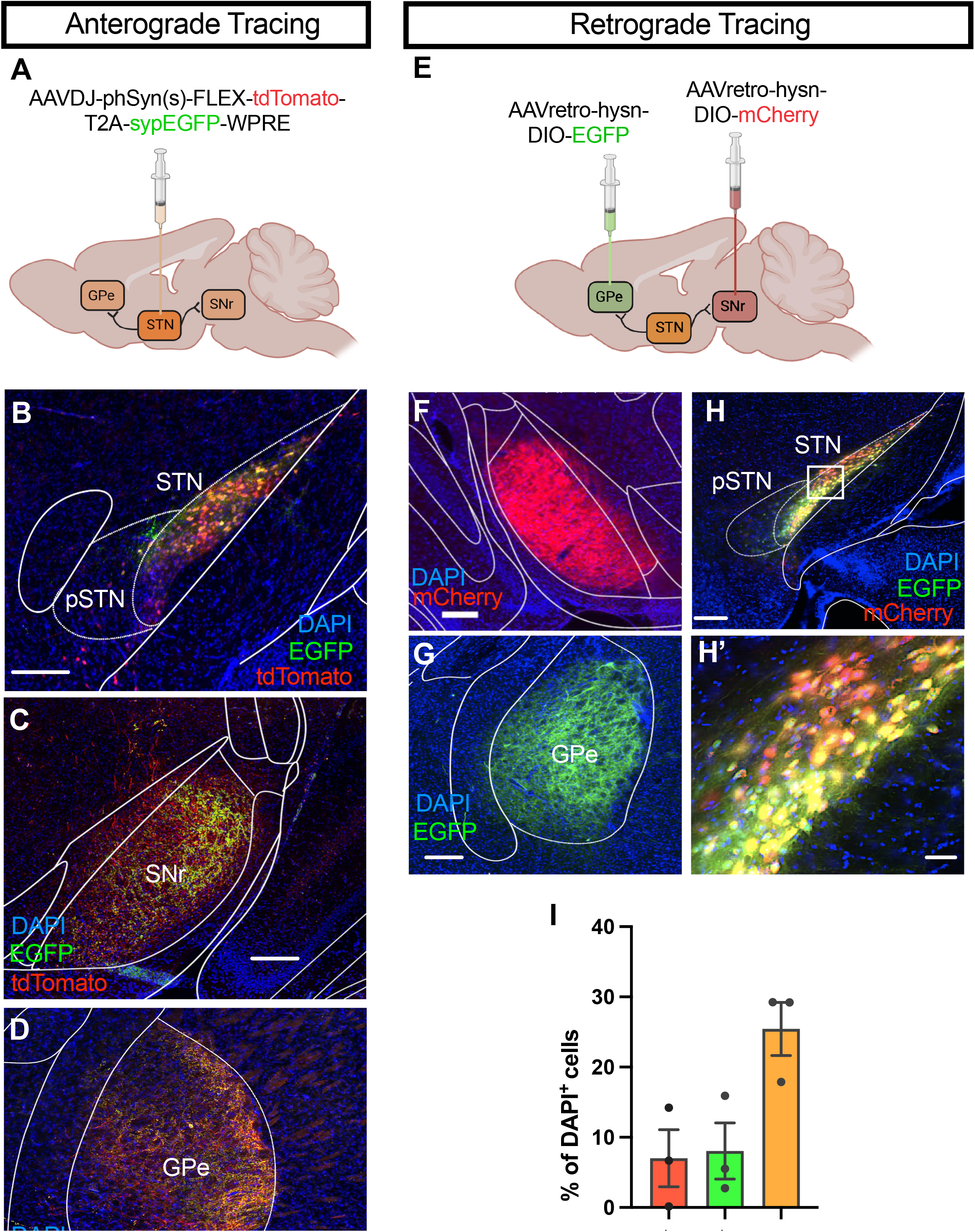
Anterograde and dual retrograde tracing reveal STN innervation of the SNr and GPe and overlapping neurons of origin. (A) Cre-dependent anterograde tracing of STN projections to the SNr and GPe (diagram was generated using BioRender.com). (B–D) Representative images of labeled STN neurons (B), STN axons (tdTomato) and synaptophysin-associated EGFP labeling in the SNr (C), and corresponding labeling in the GPe (D). (E) Dual retrograde tracing strategy using Cre-dependent EGFP and mCherry vectors injected into the GPe and SNr (diagram was generated using BioRender.com). (F,G) Representative injection sites in the SNr and GPe, respectively. (H) Retrogradely labeled STN cells; (H′) enlarged view of the boxed region in H. (I) Percentages of DAPI-positive STN cells labeled from the SNr alone, GPe alone, or both targets. Each point represents one mouse (*n* = 3); bars show group means ± SEM. Scale bars in B,C,D,F,G,H: 200 μm. Scale bar in H′: 20 μm. STN: subthalamic nucleus, pSTN: posterior subthalamic nucleus, SNr: substantia nigra pars reticulata, GPe: external globus pallidus.

### 3.5. Manipulation of STN axons in the SNr and GPe alters spontaneous motor behavior

To investigate how STN outputs contribute to spontaneous movement, we illuminated STN axons within the SNr or GPe. Cre-dependent ChR2, Jaws, or control eYFP vectors were injected bilaterally into the STN of Pitx2-Cre mice, and optical fibers were implanted above either target (Fig. 4A,B). For excitation experiments, photostimulation was delivered at 20 Hz, the frequency associated with the largest observed reduction in mobile time during somatic excitation.

**Figure 4.**
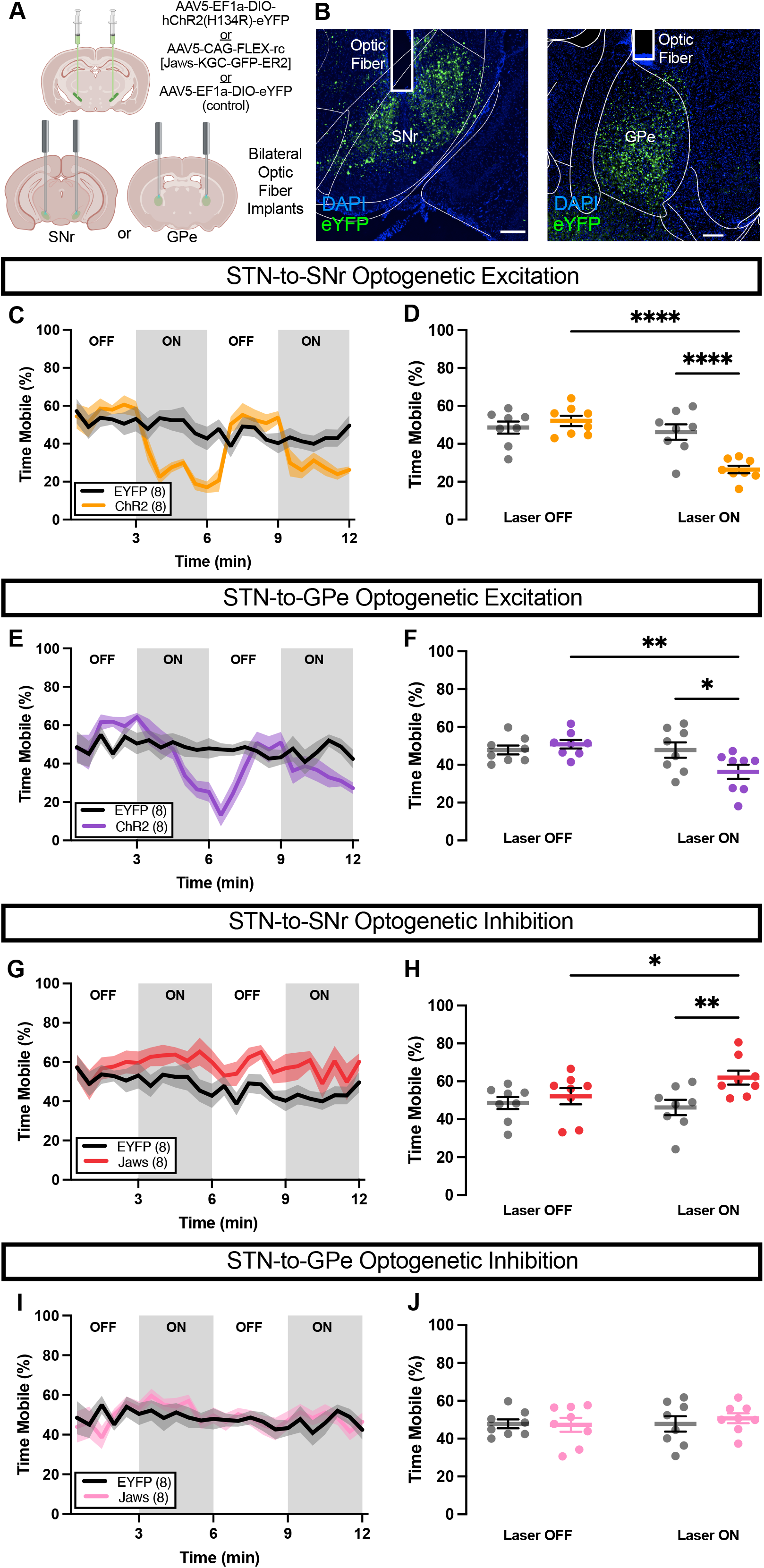
Excitation and inhibition of STN axons in the SNr and GPe alter open-field mobility. (A) Bilateral STN delivery of ChR2, Jaws, or control eYFP vectors and optical fibers above the SNr or GPe (diagram was generated using BioRender.com). (B) Representative axonal fluorescence and fiber locations in the SNr (left) and GPe (right). (C,D) Mobility time course and summary during 20-Hz SNr-targeted excitation. (E,F) GPe-targeted excitation. (G,H) Continuous SNr-targeted inhibition. (I,J) GPe-targeted inhibition. Gray shading denotes illumination periods. Data are shown as individual mice and group means ± SEM; *n* = 8 mice per virus group. Two-way repeated measures ANOVA followed by unadjusted Fisher’s LSD comparisons. Asterisks denote the bracketed pairwise comparisons: \**p* < 0.05, \*\**p* < 0.01, \*\*\**p* < 0.001, \*\*\*\**p* < 0.0001. Scale bar in B: 200 μm. SNr: susbtantia nigra pars reticulata, GPe: external globus pallidus.

Excitation of STN axons within either target reduced the percentage of time spent mobile (virus × laser interactions: SNr, *F*(1, 14) = 57.28, *p* < 0.0001; GPe, *F*(1, 14) = 5.951, *p* = 0.0286; Fig. 4C–F). Distance travelled and velocity also decreased during SNr stimulation (virus × laser interactions: distance, *F*(1, 14) = 17.06, *p* = 0.0010; velocity, *F*(1, 14) = 8.274, *p* = 0.0122; Supplementary Fig. 2A,B). Within-ChR2 comparisons indicated reductions in both measures during GPe stimulation, although neither interaction was significant (Supplementary Fig. 2D,E).

Despite these reductions, within-ChR2 comparisons indicated increased jumping during illumination within either target, although the corresponding virus × laser interactions were not significant (Supplementary Fig. 2C,F). Terminal excitation was therefore associated with intermittent jumping alongside reduced distance travelled and velocity, contrasting with the increases in these measures accompanying somatic excitation.

We next examined whether inhibition of STN axons increased spontaneous movement, as observed following somatic inhibition. In the SNr-targeted cohort, pairwise comparisons indicated increased mobile time during illumination in Jaws-expressing mice and a difference from eYFP controls during illumination, although the virus × laser interaction was not significant (*F*(1, 14) = 3.806, *p* = 0.0714; Fig. 4G,H). SNr-targeted inhibition also increased distance travelled, with a significant virus × laser interaction (*F*(1, 14) = 6.513, *p* = 0.0230; Supplementary Fig. 2G). In contrast, GPe-targeted inhibition did not significantly alter mobile time or distance travelled (Fig. 4I,J; Supplementary Fig. 2J). Velocity did not change significantly within either Jaws group (Supplementary Fig. 2H,K), and no jumping was observed (Supplementary Fig. 2I,L). Thus, SNr-targeted inhibition reproduced part of the increase in spontaneous movement observed during somatic inhibition.

### 3.6. Excitation of STN axons within the SNr or GPe disrupts rotarod performance

Finally, we assessed whether manipulating STN axons within the SNr or GPe affected motor task performance on the rotarod. At 20 Hz, photostimulation shortened latency to fall following illumination within either the SNr (virus × laser interaction, *F*(1, 14) = 2,853, *p* < 0.0001; Fig. 5A) or GPe (*F*(1, 14) = 6,688, *p* < 0.0001; Fig. 5B). As observed during somatic excitation, mice frequently jumped from the apparatus immediately after laser onset, suggesting that the reduced latency was largely driven by stimulation-induced jumping.

**Figure 5.**
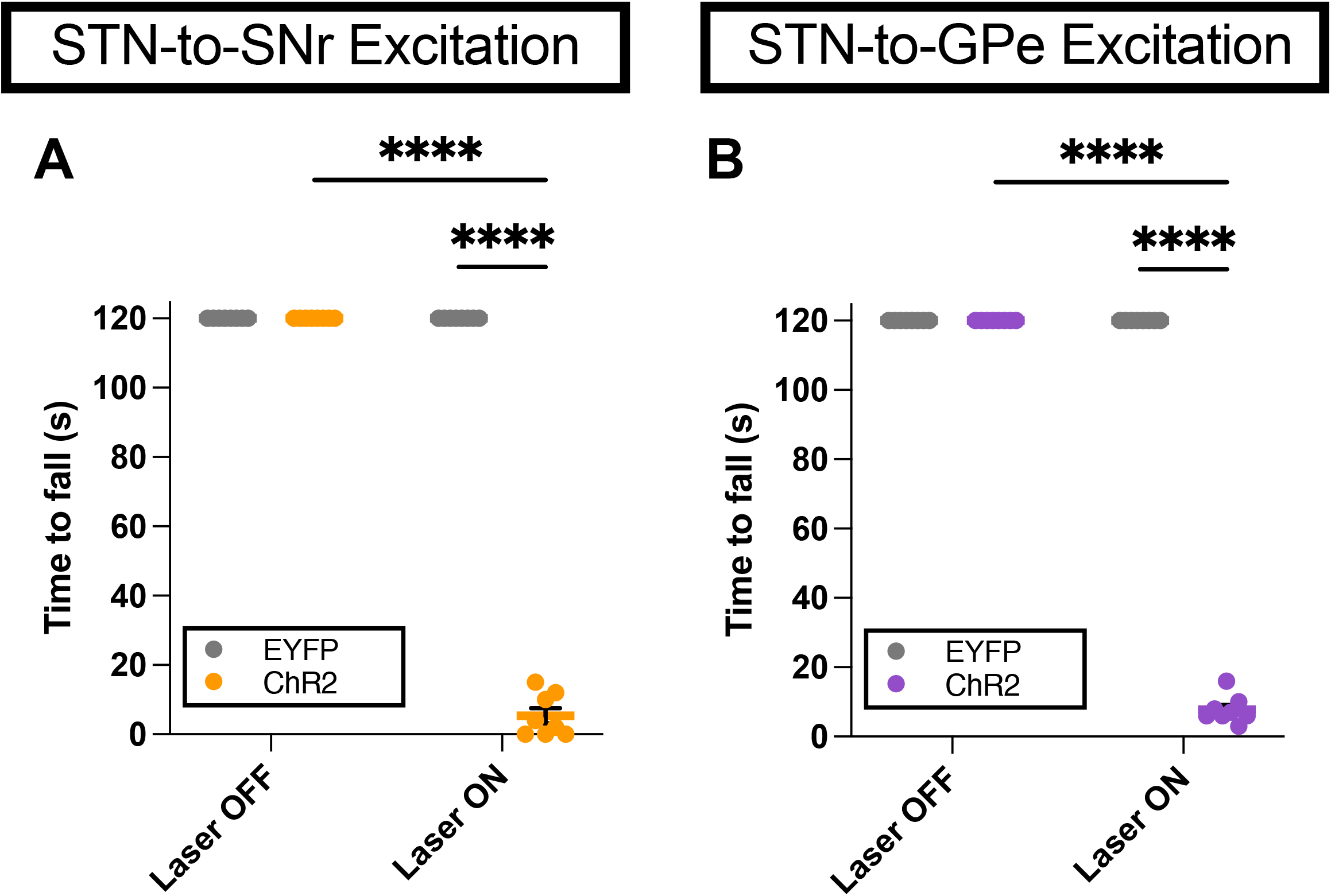
Excitation of STN axons in either the SNr or GPe shortens rotarod latency. Latency to fall during OFF and ON trials with 20-Hz excitation of STN axons in the SNr (A) or GPe (B). Data are shown as individual mice and group means ± SEM; *n* = 8 mice per virus group. Two-way repeated measures ANOVA followed by unadjusted Fisher’s LSD comparisons. Asterisks denote the bracketed pairwise comparisons: \**p* < 0.05, \*\**p* < 0.01, \*\*\**p* < 0.001, \*\*\*\**p* < 0.0001.

In contrast, inhibition of STN axons did not significantly alter rotarod latency in either target group (virus × laser interactions, both *F*(1, 14) = 1.000, *p* = 0.3343; Supplementary Fig. 3A,B). Thus, excitation within either target reproduced the disruption observed during somatic STN excitation, whereas inhibition of these projections individually did not reproduce the impairment associated with somatic inhibition.

## 4. Discussion

The STN is commonly associated with movement suppression, but its effects on motor behavior differed across somatic and axonal manipulations. Somatic excitation suppressed spontaneous mobility while eliciting jumping, whereas inhibition increased movement; both manipulations disrupted rotarod performance. Excitation of STN axons within either the SNr or GPe reproduced key features of the somatic excitation phenotype, while SNr-targeted inhibition promoted movement without a detectable change in rotarod performance. Together, these findings support a role for STN outputs in restraining spontaneous movement, while showing that changes in mobility and motor task performance were not consistently accompanied by corresponding changes in motor task performance.

### 4.1. STN activity regulates spontaneous movement beyond locomotor suppression

The reduction in mobility during STN excitation and its increase during inhibition are consistent with classical basal ganglia models, in which STN excitation of output nuclei strengthens inhibition of downstream motor circuits (Nambu et al., 2002). These findings agree with previous optogenetic experiments in Pitx2-Cre mice (Guillaumin et al., 2021) and with hyperactivity following elimination of cortico-STN hyperdirect projections (Koketsu et al., 2021). Together, these studies support the proposal that STN activity constrains spontaneous locomotion.

Excitation nevertheless elicited jumping, showing that its effects were not uniformly suppressive across behaviors. During somatic stimulation, increased distance travelled and velocity likely reflected brief jumps across the arena interspersed with periods of reduced mobility. Terminal stimulation produced a different combination: pairwise analyses indicated increased jumping alongside reduced distance and velocity. Jumping therefore did not necessarily produce a net increase in tracked displacement over the illumination period. Reduced locomotion may have outweighed displacement contributed by jumps, although movement components were not separately quantified. Differences in jump frequency or trajectory, or in the extent of locomotor suppression, could contribute to the contrast with somatic stimulation; these possibilities remain untested.

This interpretation is consistent with the jumping reported by Guillaumin et al. (2021) and with evidence that STN output can generate graded head and body movements through brainstem projections (Friedman and Yin, 2023). Our findings extend this work by suggesting that stimulation within nigral and pallidal projection territories can also elicit jumping. Whether these responses reflect recruitment of particular motor circuits or an aversive state remains unresolved. Jumping alone does not establish escape motivation, but its occurrence alongside reduced mobility illustrates why the effects of STN activation cannot be described adequately as generalized motor inhibition.

### 4.2. Motor task performance requires appropriately regulated STN activity

Both somatic excitation and inhibition impaired the ability to remain on the rotarod, despite their opposing effects on spontaneous mobility. Successful performance requires stepping and postural adjustments matched to the rotating surface. Observations of jumping during excitation and forward running during inhibition suggest that different behavioral responses contributed to reduced latency. Increased open-field movement after SNr-targeted inhibition, without a significant change in rotarod latency, further distinguishes spontaneous activity from performance on this task.

These findings are compatible with successful task performance depending on STN activity remaining within an appropriate range, although they do not establish an optimal firing rate. Suppressing the STN in monkeys increases pallidal spike-train variability and prolongs movement time (Hasegawa et al., 2022), while movement-related STN activity supports ongoing locomotion in mice (Callahan et al., 2024). Normal STN function therefore includes contributions to sustaining movement as well as restricting it, consistent with models emphasizing the temporal organization of basal ganglia signaling (Chiken et al., 2021; Nambu et al., 2023).

Our rotarod findings cannot be attributed specifically to impaired balance or coordination. Guillaumin et al. (2021) reported motor-task impairments independent of jumping, whereas jumping frequently accounted for departure from the rod in our excitation experiments. Likewise, increased mobility during somatic inhibition contrasts with the slowing following brief lateral STN inhibition reported by Callahan et al. (2024). Differences in illumination duration, anatomical coverage, and behavioral demands may contribute to these discrepancies.

### 4.3. Nigral and pallidal outputs contribute to motor control through overlapping projections

Excitation of STN axons within either the SNr or GPe reduced spontaneous mobility, supporting a contribution of both projection territories to movement restraint. SNr-targeted excitation also reduced distance travelled and velocity, while inhibition increased distance and was associated with increased mobile time. Evidence for reduced distance and velocity during GPe stimulation, increased jumping during stimulation within either target, and increased mobile time during SNr inhibition came from pairwise comparisons without significant virus × laser interactions and therefore requires more cautious interpretation.

Excitation of STN inputs to the SNr could strengthen inhibitory output to downstream motor structures (Freeze et al., 2013; Kita and Kitai, 1987). In the GPe, consequences depend on interacting cell populations: STN input can excite prototypic neurons while indirectly inhibiting arkypallidal neurons (Aristieta et al., 2021). Recordings from identified target neurons will be needed to connect these circuit responses to behavior.

Dual retrograde labeling demonstrated that individual Pitx2-Cre-targeted STN neurons supply both targets, consistent with anatomical descriptions of axonal branching (Sato et al., 2000). Labeling-category percentages did not differ significantly. While numerically smaller populations of neurons were positive for markers from a single projection target, it should be noted that single labeling does not necessarily establish exclusive projection because tracer uptake may be incomplete. Shared projections also limit claims of independent pathway control: axonal stimulation can recruit parent neurons and other collaterals through antidromic activation (Hsu et al., 2025). Similar responses to SNr and GPe illumination may therefore reflect local synaptic actions, broader recruitment of shared outputs, or both.

Although SNr-targeted inhibition reproduced part of the somatic open-field phenotype, neither terminal inhibition manipulation reproduced the rotarod impairment. This could reflect contributions from additional or combined outputs, but incomplete terminal inhibition remains an alternative. Different inhibitory opsins were used for somatic and axonal experiments as ArchT failed to express sufficiently in terminal regions, and synaptic suppression was not physiologically quantified. Given the biophysical constraints of presynaptic optogenetic inhibition (Mahn et al., 2016), these negative results do not establish that either projection is dispensable.

### 4.4. Implications for understanding STN dysfunction

The coexistence of movement suppression, jumping, and disrupted task performance is relevant to interpreting STN interventions in Parkinson’s disease, where lesions, inactivation, and deep brain stimulation can alleviate motor symptoms (Bergman et al., 1990; Krack et al., 2003; Wichmann et al., 1994). However, experiments in otherwise healthy mice do not establish the effects of these manipulations in dopamine-depleted circuits.

Nor should excitation-induced disruption be equated with therapeutic stimulation. High-frequency optogenetic STN stimulation can improve parkinsonian motor behavior under different stimulation conditions (Yu et al., 2020). Our findings motivate examination of how interventions recruit overlapping STN outputs and affect specific movements. Direct measurements of stepping, posture, and jump trajectories should help determine how stimulation alters spontaneous behavior while preserving the activity required for sustained motor performance.

## 5. Conclusion

STN excitation and inhibition had opposing effects on spontaneous mobility, yet both disrupted rotarod performance, suggesting that successful motor task performance requires appropriately regulated STN activity. Excitation within nigral and pallidal projection territories reproduced key effects of somatic excitation, whereas SNr-targeted inhibition increased spontaneous movement without detectable rotarod impairment. These findings implicate overlapping STN outputs in restraining spontaneous movement while distinguishing this function from the regulation of ongoing motor task performance.

## Lead contact

Requests for further information and resources should be directed to and will be fulfilled by the lead contact, Takatoshi Hikida.

## Materials availability

This study did not generate new unique reagents.

## CRediT authorship contribution statement

**Tom Macpherson**: Conceptualization, Investigation, Formal Analysis, Writing: Original Draft, Writing: Review & Editing, Visualization, Supervision, Funding Acquisition. **Tianchang Zhao**: Investigation, Formal Analysis, Writing: Original Draft, Writing: Review & Editing, Visualization. **Monica Natsumi Daudelin:** Investigation. **Tadaaki Nishioka**: Resources, Writing: Review & Editing. **Suthinee Attachaipanich**: Writing: Review & Editing, Supervision. **Takaaki Ozawa**: Writing: Review & Editing, Funding Acquisition. **Takatoshi Hikida**: Conceptualization, Writing: Review & Editing, Supervision, Funding Acquisition.

## Data availability

Behavioral data reported in this paper are available from the lead contact upon request.

## Declaration of generative AI and AI-assisted technologies in the manuscript preparation process

During preparation of this manuscript, the authors used Claude (Anthropic) to assist with literature searches, and manuscript drafting and revision. After using this tool, the authors reviewed and edited the content as needed and take full responsibility for the content of the published article.

---

**Figure S1.**
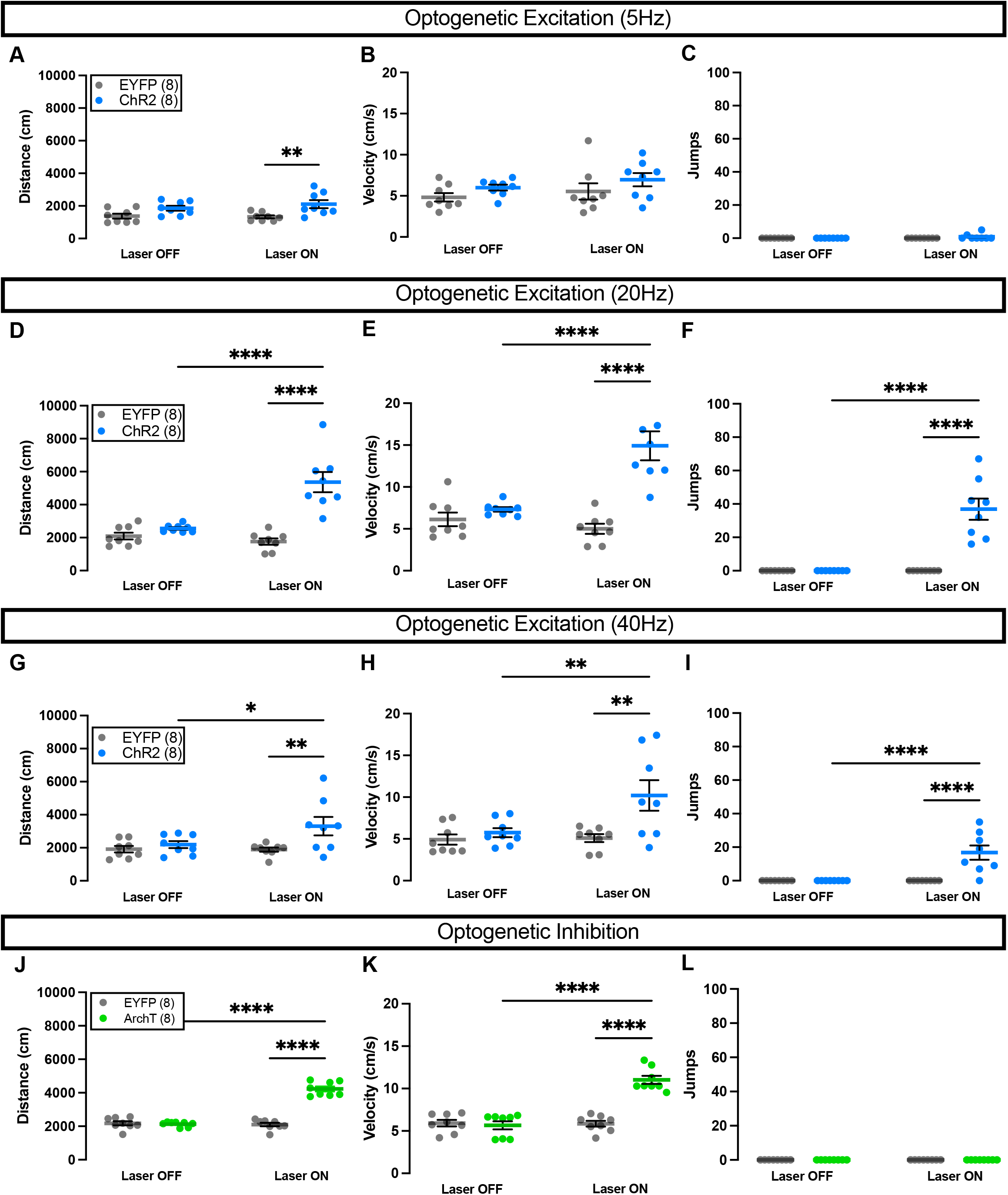
Effects of somatic STN manipulation on distance travelled, velocity, and jumping. Distance travelled (cm), velocity (cm/s), and jump count during somatic ChR2 excitation at 5 Hz (A–C), 20 Hz (D–F), and 40 Hz (G–I), and ArchT inhibition (J–L), with corresponding eYFP controls. Data are shown as individual mice and group means ± SEM; *n* = 8 mice per virus group. Two-way repeated measures ANOVA followed by unadjusted Fisher’s LSD comparisons. Asterisks denote the bracketed pairwise comparisons: \**p* < 0.05, \*\**p* < 0.01, \*\*\**p* < 0.001, \*\*\*\**p* < 0.0001.

**Figure S2.**
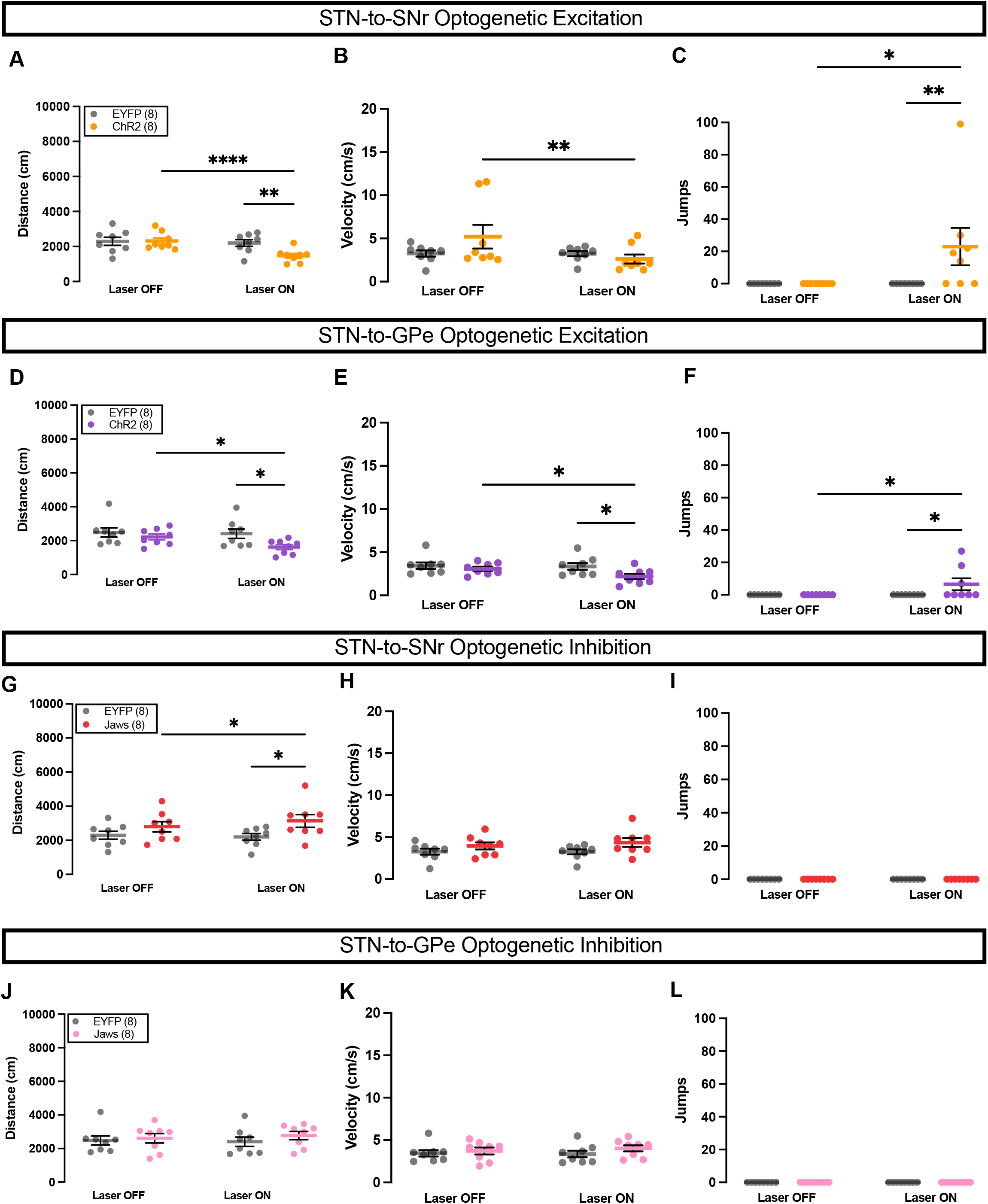
Effects of STN axonal excitation and inhibition on distance travelled, velocity, and jumping. Distance travelled (cm), velocity (cm/s), and jump count during 20-Hz excitation of STN axons in the SNr (A–C) or GPe (D–F), and continuous inhibition in the SNr (G–I) or GPe (J–L). Data are shown as individual mice and group means ± SEM; *n* = 8 mice per virus group. Two-way repeated measures ANOVA followed by unadjusted Fisher’s LSD comparisons. Asterisks denote the bracketed pairwise comparisons: \**p* < 0.05, \*\**p* < 0.01, \*\*\**p* < 0.001, \*\*\*\**p* < 0.0001.

**Figure S3.**
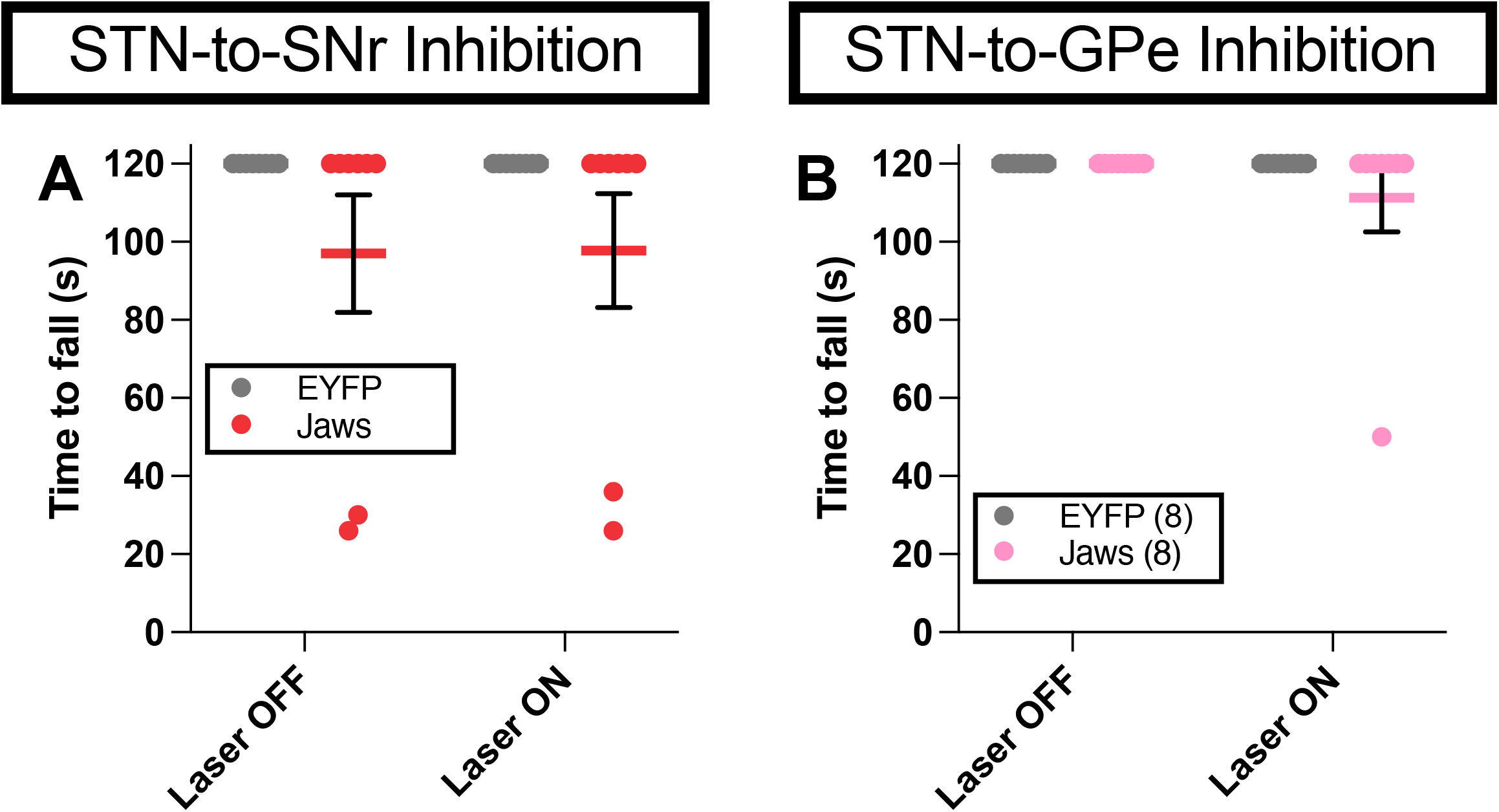
Rotarod performance during inhibition of STN axons within the SNr or GPe. Latency to fall during OFF and ON trials with continuous inhibition of STN axons in the SNr (A) or GPe (B). Data are shown as individual mice and group means ± SEM; *n* = 8 mice per virus group. Two-way repeated measures ANOVA followed by unadjusted Fisher’s LSD comparisons. Asterisks denote the bracketed pairwise comparisons: \**p* < 0.05, \*\**p* < 0.01, \*\*\**p* < 0.001, \*\*\*\**p* < 0.0001.

## References

Abdi, A., Mallet, N., Mohamed, F.Y., Sharott, A., Dodson, P.D., Nakamura, K.C., Suri, S., Avery, S.V., Larvin, J.T., Garas, F.N., Garas, S.N., Vinciati, F., Morin, S., Bezard, E., Baufreton, J., Magill, P.J., 2015. Prototypic and Arkypallidal Neurons in the Dopamine-Intact External Globus Pallidus. J. Neurosci. 35, 6667–6688. 10.1523/jneurosci.4662-14.2015

Aristieta, A., Barresi, M., Azizpour Lindi, S., Barrière, G., Courtand, G., de la Crompe, B., Guilhemsang, L., Gauthier, S., Fioramonti, S., Baufreton, J., Mallet, N.P., 2021. A Disynaptic Circuit in the Globus Pallidus Controls Locomotion Inhibition. Curr. Biol. 31, 707–721.e7. 10.1016/j.cub.2020.11.019

Attachaipanich, S., Ozawa, T., Macpherson, T., Hikida, T., 2023. Dual Roles for Nucleus Accumbens Core Dopamine D1-Expressing Neurons Projecting to the Substantia Nigra Pars Reticulata in Limbic and Motor Control in Male Mice. eNeuro 10, ENEURO.0082-23.2023. 10.1523/eneuro.0082-23.2023

Bergman, H., Wichmann, T., DeLong, M.R., 1990. Reversal of Experimental Parkinsonism by Lesions of the Subthalamic Nucleus. Science 249, 1436–1438. 10.1126/science.2402638

Callahan, J.W., Morales, J.C., Atherton, J.F., Wang, D., Kostic, S., Bevan, M.D., 2024. Movement-related increases in subthalamic activity optimize locomotion. Cell Rep. 43, 114495. 10.1016/j.celrep.2024.114495

Chiken, S., Takada, M., Nambu, A., 2021. Altered Dynamic Information Flow through the Cortico-Basal Ganglia Pathways Mediates Parkinson’s Disease Symptoms. Cereb. Cortex 31, 5363–5380. 10.1093/cercor/bhab164

Cui, G., Jun, S.B., Jin, X., Pham, M.D., Vogel, S.S., Lovinger, D.M., Costa, R.M., 2013. Concurrent activation of striatal direct and indirect pathways during action initiation. Nature 494, 238–242. 10.1038/nature11846

Fife, K.H., Gutierrez-Reed, N.A., Zell, V., Bailly, J., Lewis, C.M., Aron, A.R., Hnasko, T.S., 2017. Causal role for the subthalamic nucleus in interrupting behavior. eLife 6, e27689. 10.7554/elife.27689

Freeze, B.S., Kravitz, A.V., Hammack, N., Berke, J.D., Kreitzer, A.C., 2013. Control of Basal Ganglia Output by Direct and Indirect Pathway Projection Neurons. J. Neurosci. 33, 18531–18539. 10.1523/jneurosci.1278-13.2013

Friedman, A.D., Yin, H.H., 2023. Selective Activation of Subthalamic Nucleus Output Quantitatively Scales Movements. J. Neurosci. 43, 7967–7981. 10.1523/jneurosci.0734-23.2023

Guillaumin, A., Serra, G.P., Georges, F., Wallén-Mackenzie, Å., 2021. Experimental investigation into the role of the subthalamic nucleus (STN) in motor control using optogenetics in mice. Brain Res. 1755, 147226. 10.1016/j.brainres.2020.147226

Hasegawa, T., Chiken, S., Kobayashi, K., Nambu, A., 2022. Subthalamic nucleus stabilizes movements by reducing neural spike variability in monkey basal ganglia. Nat. Commun. 13, 2233. 10.1038/s41467-022-29750-2

Hsu, L.M., Cerri, D.H., Carelli, R.M., Shih, Y.Y.I., 2025. Optogenetic stimulation of cell bodies versus axonal terminals generate comparable activity and functional connectivity patterns in the brain. Brain Stimul. 18, 822–828. 10.1016/j.brs.2025.03.006

Isoda, M., Hikosaka, O., 2008. Role for Subthalamic Nucleus Neurons in Switching from Automatic to Controlled Eye Movement. J. Neurosci. 28, 7209–7218. 10.1523/jneurosci.0487-08.2008

Ji, Y.W., Zhang, X., Fan, J.P., Gu, W.X., Shen, Z.L., Wu, H.C., Cui, G., Zhou, C., Xiao, C., 2023. Differential remodeling of subthalamic projections to basal ganglia output nuclei and locomotor deficits in 6-OHDA-induced hemiparkinsonian mice. Cell Rep. 42, 112178. 10.1016/j.celrep.2023.112178

Kita, H., Kitai, S.T., 1987. Efferent projections of the subthalamic nucleus in the rat: Light and electron microscopic analysis with the PHA-L method. J. Comp. Neurol. 260, 435–452. 10.1002/cne.902600309

Koketsu, D., Chiken, S., Hisatsune, T., Miyachi, S., Nambu, A., 2021. Elimination of the Cortico-Subthalamic Hyperdirect Pathway Induces Motor Hyperactivity in Mice. J. Neurosci. 41, 5502–5510. 10.1523/jneurosci.1330-20.2021

Krack, P., Batir, A., Van Blercom, N., Chabardes, S., Fraix, V., Ardouin, C., Koudsie, A., Limousin, P.D., Benazzouz, A., LeBas, J.F., Benabid, A.L., Pollak, P., 2003. Five-Year Follow-up of Bilateral Stimulation of the Subthalamic Nucleus in Advanced Parkinson’s Disease. N. Engl. J. Med. 349, 1925–1934. 10.1056/nejmoa035275

Kravitz, A.V., Freeze, B.S., Parker, P.R.L., Kay, K., Thwin, M.T., Deisseroth, K., Kreitzer, A.C., 2010. Regulation of parkinsonian motor behaviours by optogenetic control of basal ganglia circuitry. Nature 466, 622–626. 10.1038/nature09159

Macpherson, T., Morita, M., Hikida, T., 2014. Striatal direct and indirect pathways control decision-making behavior. Front. Psychol. 5, 1301. 10.3389/fpsyg.2014.01301.

Macpherson, T., Matsumoto, M., Gomi, H., Morimoto, J., Uchibe, E., Hikida, T., 2021. Parallel and hierarchical neural mechanisms for adaptive and predictive behavioral control. Neural Netw. 144, 507–521. 10.1016/j.neunet.2021.09.009.

Mahn, M., Prigge, M., Ron, S., Levy, R., Yizhar, O., 2016. Biophysical constraints of optogenetic inhibition at presynaptic terminals. Nat. Neurosci. 19, 554–556. 10.1038/nn.4266

Mallet, N., Micklem, B.R., Henny, P., Brown, M.T., Williams, C., Bolam, J.P., Nakamura, K.C., Magill, P.J., 2012. Dichotomous Organization of the External Globus Pallidus. Neuron 74, 1075–1086. 10.1016/j.neuron.2012.04.027

Martin, D.M., Skidmore, J.M., Philips, S.T., Vieira, C., Gage, P.J., Condie, B.G., Raphael, Y., Martinez, S., Camper, S.A., 2004. PITX2 is required for normal development of neurons in the mouse subthalamic nucleus and midbrain. Dev. Biol. 267, 93–108. 10.1016/j.ydbio.2003.10.035

Nambu, A., Takada, M., Inase, M., Tokuno, H., 1996. Dual somatotopical representations in the primate subthalamic nucleus: evidence for ordered but reversed body-map transformations from the primary motor cortex and the supplementary motor area. J. Neurosci. 16, 2671–2683. 10.1523/jneurosci.16-08-02671.1996

Nambu, A., Tokuno, H., Hamada, I., Kita, H., Imanishi, M., Akazawa, T., Ikeuchi, Y., Hasegawa, N., 2000. Excitatory Cortical Inputs to Pallidal Neurons Via the Subthalamic Nucleus in the Monkey. J. Neurophysiol. 84, 289–300. 10.1152/jn.2000.84.1.289

Nambu, A., Tokuno, H., Takada, M., 2002. Functional significance of the cortico–subthalamo–pallidal ‘hyperdirect’ pathway. Neurosci. Res. 43, 111–117. 10.1016/s0168-0102(02)00027-5

Nambu, A., Chiken, S., Sano, H., Hatanaka, N., Obeso, J.A., 2023. Dynamic Activity Model of Movement Disorders: The Fundamental Role of the Hyperdirect Pathway. Mov. Disord. 38, 2145–2150. 10.1002/mds.29646

Nishioka, T., Attachaipanich, S., Hamaguchi, K., Lazarus, M., de Kerchove d’Exaerde, A., Macpherson, T., Hikida, T., 2023. Error-related signaling in nucleus accumbens D2 receptor-expressing neurons guides inhibition-based choice behavior in mice. Nat. Commun. 14, 2284. 10.1038/s41467-023-38025-3.

Nishioka, T., Hamaguchi, K., Yawata, S., Hikida, T., Watanabe, D., 2020. Chemogenetic Suppression of the Subthalamic Nucleus Induces Attentional Deficits and Impulsive Action in a Five-Choice Serial Reaction Time Task in Mice. Front. Syst. Neurosci. 14, 38. 10.3389/fnsys.2020.00038

Parolari, L., Schneeberger, M., Heintz, N., Friedman, J.M., 2021. Functional analysis of distinct populations of subthalamic nucleus neurons on Parkinson’s disease and OCD-like behaviors in mice. Mol. Psychiatry 26, 7029–7046. 10.1038/s41380-021-01162-6

Polyakova, Z., Chiken, S., Hatanaka, N., Nambu, A., 2020. Cortical Control of Subthalamic Neuronal Activity through the Hyperdirect and Indirect Pathways in Monkeys. J. Neurosci. 40, 7451–7463. 10.1523/jneurosci.0772-20.2020

Polyakova, Z., Hatanaka, N., Chiken, S., Nambu, A., 2024. Subthalamic Activity for Motor Execution and Cancellation in Monkeys. J. Neurosci. 44, e1911222024. 10.1523/jneurosci.1911-22.2024

Roseberry, T.K., Lee, A.M., Lalive, A.L., Wilbrecht, L., Bonci, A., Kreitzer, A.C., 2016. Cell-Type-Specific Control of Brainstem Locomotor Circuits by Basal Ganglia. Cell 164, 526–537. 10.1016/j.cell.2015.12.037

Sano, H., Chiken, S., Hikida, T., Kobayashi, K., Nambu, A., 2013. Signals through the Striatopallidal Indirect Pathway Stop Movements by Phasic Excitation in the Substantia Nigra. J. Neurosci. 33, 7583–7594. 10.1523/jneurosci.4932-12.2013

Sato, F., Parent, M., Levesque, M., Parent, A., 2000. Axonal branching pattern of neurons of the subthalamic nucleus in primates. J. Comp. Neurol. 424, 142–152. https://doi.org/10.1002/1096-9861(20000814)424:1<142::aid-cne10>3.0.co;2-8

Schmidt, R., Leventhal, D.K., Mallet, N., Chen, F., Berke, J.D., 2013. Canceling actions involves a race between basal ganglia pathways. Nat. Neurosci. 16, 1118–1124. 10.1038/nn.3456

Schweizer, N., Pupe, S., Arvidsson, E., Nordenankar, K., Smith-Anttila, C.J., Mahmoudi, S., Andrén, A., Dumas, S., Rajagopalan, A., Lévesque, D., Leão, R.N., Wallén-Mackenzie, Å., 2014. Limiting glutamate transmission in a Vglut2-expressing subpopulation of the subthalamic nucleus is sufficient to cause hyperlocomotion. Proc. Natl. Acad. Sci. U.S.A. 111, 7837–7842. 10.1073/pnas.1323499111

Schweizer, N., Viereckel, T., Smith-Anttila, C.J., Nordenankar, K., Arvidsson, E., Mahmoudi, S., Zampera, A., Wärner Jonsson, H., Bergquist, J., Lévesque, D., Konradsson-Geuken, Å., Andersson, M., Dumas, S., Wallén-Mackenzie, Å., 2016. Reduced Vglut2/Slc17a6 Gene Expression Levels throughout the Mouse Subthalamic Nucleus Cause Cell Loss and Structural Disorganization Followed by Increased Motor Activity and Decreased Sugar Consumption. eNeuro 3, ENEURO.0264-16.2016. 10.1523/ENEURO.0264-16.2016

Takakusaki, K., Habaguchi, T., Ohtinata-Sugimoto, J., Saitoh, K., Sakamoto, T., 2003. Basal ganglia efferents to the brainstem centers controlling postural muscle tone and locomotion: a new concept for understanding motor disorders in basal ganglia dysfunction. Neuroscience 119, 293–308. 10.1016/s0306-4522(03)00095-2

Tye, K.M., Mirzabekov, J.J., Warden, M.R., Ferenczi, E.A., Tsai, H.C., Finkelstein, J., Kim, S.Y., Adhikari, A., Thompson, K.R., Andalman, A.S., Gunaydin, L.A., Witten, I.B., Deisseroth, K., 2013. Dopamine neurons modulate neural encoding and expression of depression-related behaviour. Nature 493, 537–541. 10.1038/nature11740

Wallén-Mackenzie, Å., Dumas, S., Papathanou, M., Martis Thiele, M.M., Vlcek, B., König, N., Björklund, Å.K., 2020. Spatio-molecular domains identified in the mouse subthalamic nucleus and neighboring glutamatergic and GABAergic brain structures. Commun. Biol. 3, 338. 10.1038/s42003-020-1028-8

Watson, G.D.R., Hughes, R.N., Petter, E.A., Fallon, I.P., Kim, N., Severino, F.P.U., Yin, H.H., 2021. Thalamic projections to the subthalamic nucleus contribute to movement initiation and rescue of parkinsonian symptoms. Sci. Adv. 7, eabe9192. 10.1126/sciadv.abe9192

Wichmann, T., Bergman, H., DeLong, M.R., 1994. The primate subthalamic nucleus. III. Changes in motor behavior and neuronal activity in the internal pallidum induced by subthalamic inactivation in the MPTP model of parkinsonism. J. Neurophysiol. 72, 521–530. 10.1152/jn.1994.72.2.521

Yu, C., Cassar, I.R., Sambangi, J., Grill, W.M., 2020. Frequency-Specific Optogenetic Deep Brain Stimulation of Subthalamic Nucleus Improves Parkinsonian Motor Behaviors. J. Neurosci. 40, 4323–4334. 10.1523/jneurosci.3071-19.2020

Zhou, J., Sajid, M.S., Hormigo, S., Castro-Alamancos, M.A., 2026. The Subthalamic Nucleus Controls Action Timing under Threat. eNeuro 13, ENEURO.0065-26.2026. 10.1523/eneuro.0065-26.2026

